# Instantaneous Brain Dynamics Mapped to a Continuous State Space

**DOI:** 10.1101/157115

**Authors:** Jacob Billings, Alessio Medda, Sadia Shakil, Xiaohong Shen, Amrit Kashyap, Shiyang Chen, Anzar Abbas, Xiaodi Zhang, Maysam Nezafati, Wen-Ju Pan, Gordon Berman, Shella Keilholz

**Affiliations:** Emory University, Graduate Division of Biological and Biomedical Sciences – Program in Neuroscience, Atlanta, USA; Georgia Tech Research Institute, Division of Aerospace & Acoustics Technologies, Atlanta, USA; Georgia Institute of Technology, School of Electrical and Computer Engineering, Atlanta, USA; Institute of Space Technology, Department of Electrical Engineering, Islamabad, Pakistan; Georgia Institute of Technology and Emory University, Department of Biomedical Engineering, Atlanta, USA; Shandong University of Finance and Economics of China, School of Computer Science and Technology, Jinan, China; Georgia Institute of Technology, School of Biological Engineering, Atlanta, USA; Emory University, Department of Biology, Atlanta, USA

**Keywords:** fMRI, connectivity dynamics, functional connectivity, multiscale, dimensionality reduction

## Abstract

Measures of whole-brain activity, from techniques such as functional Magnetic Resonance Imaging, provide a means to observe the brain’s dynamical operations. However, interpretation of whole-brain dynamics has been stymied by the inherently high-dimensional structure of brain activity. The present research addresses this challenge through a series of scale transformations in the spectral, spatial, and relational domains. Instantaneous multispectral dynamics are first developed from input data via a wavelet filter bank. Voxel-level signals are then projected onto a representative set of spatially independent components. The correlation distance over the instantaneous wavelet-ICA state vectors is a graph that may be embedded onto a lower-dimensional space to assist the interpretation of state-space dynamics. Applying this procedure to a large sample of resting and task data (acquired through the Human Connectome Project), we segment the empirical state space into a continuum of stimulus-dependent brain states. We also demonstrate that resting brain activity includes brain states that are very similar to those adopted during some tasks, as well as brain states that are distinct from experimentally-defined tasks. Back-projection of segmented brain states onto the brain’s surface reveals the patterns of brain activity that support each experimental state.

## Highlights

- We demonstrate the construction and interrogation of a continuous, two-dimensional map of fMRI dynamics.
- Map points represent an individual’s multispectral, and multispectral BOLD state centered at a single point in time.
- Task-based scans occupy focal state-spaces, reinforcing the utility of study methods to capture salient BOLD dynamics evoked by experimental stimuli.
- Resting-state scans occupy a broad state-space, reinforcing the view that the resting mind is highly active.

## 1. Introduction

The advent of functional Magnetic Resonance Imaging (fMRI) has launched the brain sciences into an exciting frontier by allowing the direct observation of systems-wide activity from healthy human brains (Rosen and Savoy, 2012). The richness of data this technology generates is the subject of cutting-edge research to interpret spontaneous signal fluctuations as indicators of preferential information exchange among the brain’s intrinsic networks—i.e., its functional connectivity (FC) (Biswal et al., 1995; Hutchison et al., 2013). Brain FC networks were first defined over relatively long periods of time. Such *static* FC studies reveal that brain FC naturally develops a small-world topology, where densely connected local modules communicate with one another via richly interconnected hubs (Achard et al., 2006; Bullmore and Sporns, 2009). But the brain is not a static system. Rather, differential information exchange among neurons, circuits, and networks enable brains to deal flexibly with ever-changing environmental stimuli. The availability of rapid (< 1s), whole-brain imaging prompted researchers to look for shorter term *dynamics* of brain FC (Deco et al., 2011).

Early efforts to characterize brain dynamics observed that intra-network membership and inter–network communication possessed statistically significant differences when samples were drawn from short time windows during various epochs of an fMRI scan (Chang and Glover, 2010; Keilholz et al., 2013; Smith et al., 2012; Zalesky et al., 2014). While these short time window studies confirmed the expectation that the Blood-Oxygen Level Dependent (BOLD) fMRI signal may convey information about short-term brain-state dynamics, the large effect that *a priori* choices in window length had on study results lessened the method’s analytic utility (Shakil et al., 2016). The effort to identify rapidly changing dynamics is also hampered by the drop-off in bold SNR at short window lengths.

To avoid the problems inherent in windowed analysis techniques, we present a method that provides a 2D map of the relative similarity of the brain’s activity for all time points in the scan. The signal from each voxel first undergoes wavelet decomposition, making use of the BOLD signal’s natural spectral scaling to characterize each time point as a summation of activations at multiple frequencies (Billings et al., 2015; Chang and Glover, 2010; Yaesoubi et al., 2015). This multispectral interpretation has been suggested to provide a parsimonious representation of the dynamic properties of complex systems like brains (Bullmore et al., 2004; Ciuciu et al., 2012; Mallat, 1989; Mandelbrot, 1983). To reduce the redundancy of spatial information and improve the SNR, voxel-wise signals are aggregated into a lower–dimensional spatial parcellation using Independent Component Analysis (ICA). In the present study, we treat the collected vectors of multispectral activations from all of the ICA networks at each time point as samples of instantaneous brain states.

The dimensionality of the resulting data set is high (equal to the product of the number of functional networks and the number of spectral filters) and difficult to interpret. In order to explore the dynamics of brain activity, we apply t-distributed stochastic neighbor embedding (t-SNE) to represent the data from each time point in a two dimensional space (van der Maaten and Hinton, 2008), using correlation as a distance measure to ensure that similar states are grouped together. t-SNE is a state of the art data-driven dimensionality reduction algorithm that maintains local distance structure and has found wide application in the data-driven sciences to produce visualizations of *drosophila* behavior, machine learning hidden layers, static functional connectivity networks, and a host of other multidimensional structures (Berman et al., 2014; Mnih et al., 2015; Plis et al., 2014). In comparison to clustering based approaches that segment the time course into a number of predefined states, the map created by t-SNE produces a continuous distribution that can then be segmented empirically (using the watershed algorithm in this study). Information about the timing and the relative similarity of different states is preserved.

Towards the goal of detailing a map of brain-state dynamics, the present study analyzes the wide-ranging states 446 normal volunteers adopt as part of the Human Connectome Project (HCP)(Van Essen et al., 2012b). BOLD fMRI scans from 7 distinct tasks (EMOTION, GAMBLING, LANGUAGE, MOTOR, RELATIONAL, SOCIAL, and WORKING MEMORY (WM)), and from repeated resting conditions (REST1, and REST2) provide a basis to segment a t-SNE embedding of brain-state dynamics across experimentally defined events. We demonstrate the utility of the t-SNE mapping to characterize the human brain’s coordination across time, space, and spectra during rest and in the negotiation of changing experimental stimuli.

## 2. Methods

*Data Acquisition and Preprocessing*. The data for this study was obtained from the HCP (Van Essen et al., 2012b). Whole-brain, BOLD-weighted, gradient-echo EPI data were acquired with a *TR* = 0.720 ms, and 2.0 mm isotropic voxels. Volunteers were scanned under 9 conditions, including: REST, EMOTION, GAMBLING, LANGUAGE, MOTOR, RELATIONAL, SOCIAL, and WORKING MEMORY (WM). The SOCIAL scan was examined in more detail during our analysis and is briefly described as follows: volunteers were presented 5 rounds of 20 s movies showing abstract objects making either random motions *(random)* or engaging in socially relevant movements *(mentalizing)*. Each movie is followed by a 15 s fixation period where volunteers are asked to fix look at a ‘+’ symbol. Each scan was performed twice.

A total of 446 volunteer datasets were included in the present study. Minimal data preprocessing was performed by HCP researchers. Steps included: spatial artifact and distortion removal, surface generation, anatomical registration, and alignment to grayordinate space. Voxel time series were normalized to zero mean and unit variance to fit the isotropic noise model expected by the ICA spatial filters. Each volunteer’s fMRI data was concatenated, across time, into a single matrix to minimize edge effects from spectral filtering. Scan order was randomized across volunteers.

*Analysis*. Previous studies have suggested that static functional connectivity networks segment into multiple frequency-specific architectures (Billings et al., 2015; Chang and Glover, 2010; Yaesoubi et al., 2015). Therefore, concatenated fMRI datasets were spectrally filtered into an octave of spectral bands, log-spaced over the low-frequency fluctuation range (0.1 to 0.01 Hz). Using the continuous wavelet transform schema, the filterbank was constructed from a low-order wavelet (Daubechies 4-tap wavelet) to provide optimal segmentation in the time domain with full coverage of the frequency domain (Daubechies, 1992). Brain images from each spectral band were multiplied by a 50 component group ICA spatial decomposition matrix. ICA filters were calculated as part of the HCP beta-release of group-ICA maps (Human Connectome Project, 2014). The number of components was chosen to just exceed the number needed for the eigenvalues of real and randomly shuffled data to be equal (data not shown). Time points were thus modeled as 400-dimensional states (8 spectral bands by 50 functional networks).

The state vectors for each time point were compared, pairwise, using the Pearson correlation distance. This choice highlights coordinated deviations from mean values. The correlation graph was then injected onto a 2-dimensional Euclidean surface using t-SNE. The t-SNE algorithm proceeds in two steps: first, the local neighborhood of each node is emphasized by normalizing inter-node distances via an adaptive Gaussian filter. Second, a 2-dimensional Euclidean version of the graph is constructed by minimizing the KL-divergence between the high-dimensional stochastic distribution and the low-dimensional stochastic distribution. A key innovation to t-SNE is to utilize a heavier tail in the low-dimensional probability distribution—a t-distribution rather than a Gaussian distribution. Doing so causes points that are only moderately far away in the high-dimensional space are pushed further apart in the low-dimensional representation. This feature allows for naturally affiliative clusters to emerge from an otherwise more compressed state space. Inherent similarities among sequentially sampled points will, none-the-less, cause these points to form their own distinct neighborhood. One way to encourage piecemeal-sequential points to arrive a group-level neighborhood is to embed points individually onto a group-level training embedding constructed from a sparse subsampling of each individual’s scan data. We generated the training embedding in three steps. First, concatenated time series from each volunteer’s full set of scans were t-SNE embedded into their own low-dimensional space. Second, ~2% (200 points) were selected from each volunteer’s map to construct a group-level subsample. Third, the group-level subsample was t-SNE embedded to construct the training embedding. Subsequently, out-of-sample time points were injected onto the training embedding by satisfying the same symmetrized KL-divergence as used in to generate the training embedding. This subsampling procedure has the added benefit of reducing the computational load of embedding a large number of data points. (For additional details, please see van der Maaten and Hinton (2008), Berman and Shaevitz (2014), and the supplemental materials).

*Quantitative Interpretation Methods*. Density maps were constructed by convolving embedded point distributions by a 2-dimensional Gaussian filter. Density maps were compared using the structural similarity index (SSIM). SSIM is a robust measure of inter-image similarity as it is a product of three separate terms to account for salient image features, luminance, variance, and structure (Zhou et al., 2004). SSIM values range between 0 and 1, with 1 denoting identical images. Bootstrapped distributions of these comparisons provided sample statistics for ANOVA and multiple comparisons testing (see the section entitled, “Comparing Embeddings,” in the supplemental material).

To characterize the t-SNE embedding as disjoint brain states, the watershed transform of an (inverted) density map was used to establish boundaries around peaks (the catchment basins of the inverted image) (Meyer, 1994). The granularity of the watershed map is easily adjusted by changing the width of the Gaussian curve used to make the density maps. Given a discrete segmentation of brain states into watershed regions, a rudimentary count of the amount of time participants dwell in any given region is a measure of the stability of the regionally-bounded brain state. The map space is also amenable to group-level statistical testing. For example, we constructed a null model over the hypothesis that particular experimental conditions do not have a preference for any map region. Thereby, we investigated the preference that experimentally defined conditions have for the population of brain states embedded into each map region. Finally, brain state data were projected onto the midthickness Conte69 surface-based atlas to visualize anatomical characteristics (Van Essen et al., 2012a).

*Additional methodological details, including the relevant equations, may be found in the supplemental information*.

## 3. Results

To test the degree to which resting and tasked brains develop distinct dynamics, we segmented time points during the REST1 and REST2 scans from all task-scan time points. The results are displayed as density maps in part A of figure 1. The resting brain tends to adopt a range of states in the map’s periphery, while the task-active brain tends to develop brain states at the map’s interior. To represent the brain’s dynamic transitions across the embedded state space, part B displays point-to-point state changes as a velocity field. The results demonstrate that the resting brain’s most rapid transitions occur in regions densely populated during tasks. In the task-active segmentation, the highest velocities are found at the map’s center, between two interior regions densely populated during tasks. Regions of low velocity are distributed in patches throughout the task segmentation.

**Figure 1.**
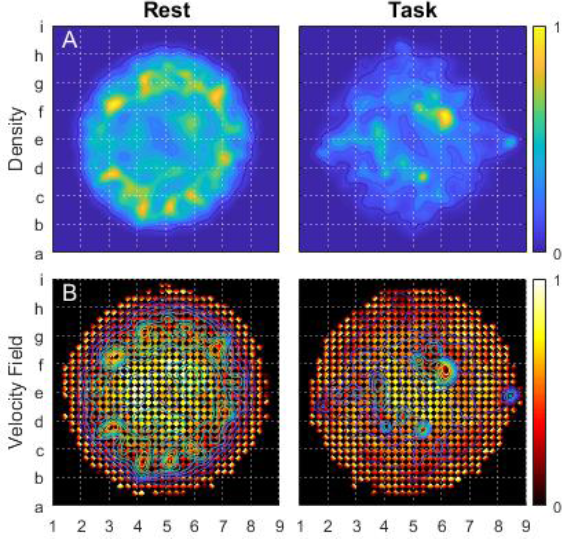
compares 2D Euclidean embeddings of instantaneous brain states, separated for the resting state and the task-active state. For ease of reference, the embedding space is divided into an 8×8 grid. Alphanumeric labels mark grid vertices. Part A displays the distribution of embedded points as a Gaussian cloud. The Gaussian filter radius equaled 1/32 the maximum point displacement from the embedding’s center. Part B displays the velocity field from an aggregation of points within a 32×32 grid in each of the 4 cardinal directions across the map space. All results were normalized to unit magnitude.

While the embedding is a continuous state space, the presence of multiple densely populated regions suggests an ensemble of discrete states that the brain adopts. Figure 2 takes on a discretized perspective by tracing boundaries around the resting-state segmentation’s dense regions. Formally, each state-space parcel is a catchment basin formed by taking the watershed transform of the density map’s inverse (Meyer, 1994). Part A of the figure codes regions in terms of the percentage of points in each parcel. Owing to its sheer size, a sprawling domain in the map’s interior contains the largest proportion of samples (4% to 5%, magenta boarder). The brain’s propensity for adopting configurations within this parcel increases during task scans, when 6% to 7% of points form a similar density (see supplemental figure S1). The average brain state within this region sustains relatively slow (~0.019 Hz), low-amplitude, in-phase activations across most of the brain’s static networks (part A, right) (see supplemental figure S2 for a description of each network). The region is often populated during the fixation periods of most tasks (see supplemental movies).

**Figure 2.**
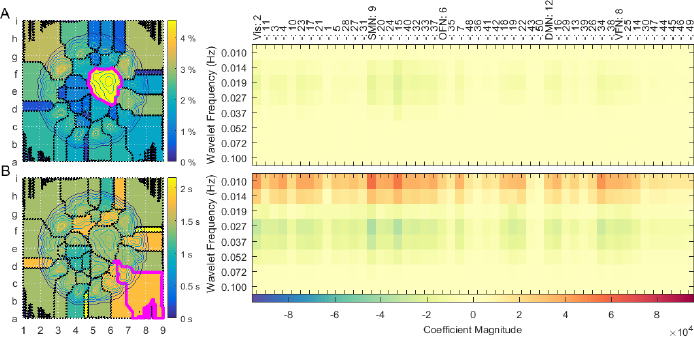
highlights the brain states adopted within watershed regions of the embedding space as participants reside in the resting-state. Part A (left) displays the percentage of points lying within each region. Part B (left) displays the median amount of time participants dwelled in each region. A similar analysis is performed for the task data in supplemental figure S1. The mean spatio-spectral brain state from the regions highlighted in magenta (left) are charted to the right. Each of the 50 ICA resting-state networks are categorized into one of five classes: ’Vis,’ visual network; ’SMN,’ somatomotor network; ’OFN,’ orbito-frontal network; ’VFN,’ ventral-frontal network.

To gain a better understanding of the dynamic characteristics of resting-state parcels, part B of figure 2 displays the median time volunteers continuously dwelt in each parcel. Although maximum resting-state dwell times reached as high as 30 s (see supplemental figure S3), median dwell times did not exceed 2.5 s during rest. This finding is a confirmation that the resting brain often transitions between states. Dwell times tended to increase in duration during tasks (see supplemental figure S1). The mean brain state of one region having a long dwell time (magenta boarder) shows the brain to sustain activations in the same ICA networks as the state highlighted in part A. However, the second region’s activations increase in magnitude. Further, they occur in two separate frequency bands, with either band’s activations flipped to the opposite phase from the other.

The wide range of experimental states adopted during HCP scans provides a natural means to segment instantaneous brain-states. Figure 3 displays group-level map densities produced by such a segmentation. Bootstrap sampling provided a sample distribution to assess structural image similarity within-states, and also between-states (figure 4). Larger within-scan SSIM values indicate that the brain adopts a tighter range of states during the scan. Comparatively large between-scan SSIM values indicate that the two scans evoke similar varieties of brain states. Multiple comparisons statistics performed on these results determined that the repeated resting-state scans, alone, bear statistically similar continuous state distributions (see supplemental figure S4). supplemental figure S5 displays the results of the same analysis when data are segmented against all block-design contrasts and task-related events.

**Figure 3.**
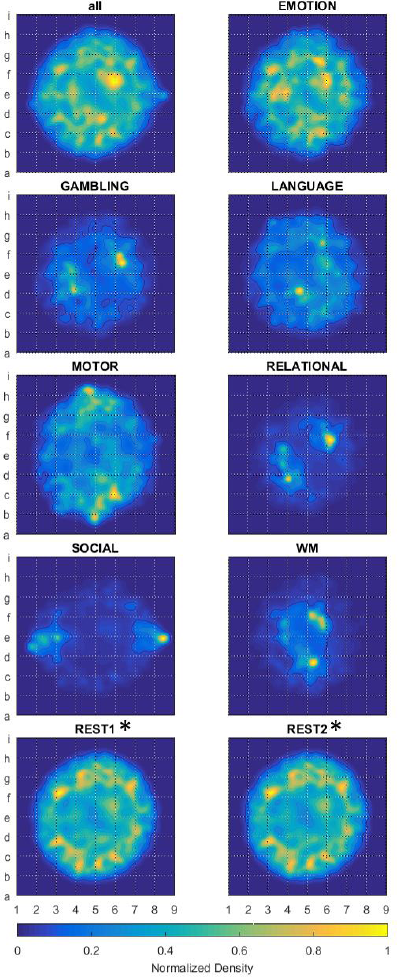
displays t-SNE embeddings of instantaneous brain states, segmented by scan, and represented as the normalized density of each scan’s embedded points. Task datasets include BOLD images during all periods of the scan, including any cue events, all contrasting task stimuli, any responses from volunteers, and any fixation blocks. The Gaussian filter radius equaled 1/32 the maximum point displacement from the embedding’s center

**Figure 4.**
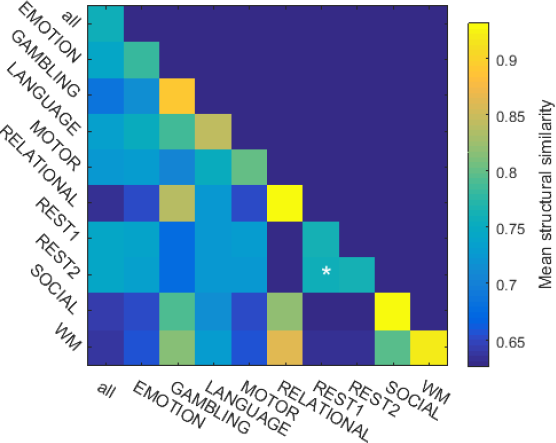
plots the mean structural similarity index (SSIM) between normalized density embeddings, segmented across scans. For each scan type, the sample distribution was bootstrapped from 50 realizations of 2500 timepoints, randomly sampled from the group-level data set. The number of time points provides a representative sampling of each scan’s embedded distribution. Asterisks indicate between-scan comparisons whose mean SSIM was not significantly less than either within-scan mean SSIM.

The block design of HCP task scans—where stimuli are presented in rigidly timed sequences over several blocks—makes it possible to identify significant differences in the point-wise evolution of brain-states during the navigation of contrasting tasks, i.e., fine-scale brain dynamics. While the main text of the present manuscript uses the SOCIAL scan as an example, similar results are found for each set of block-design contrasts. (Movies illustrating the time-locked brain state distributions for all block-design contrasts may be found in the supplemental materials). Figure 5 outlines the point-wise state transitions during each SOCIAL task contrast’s 35 s block. A bar chart of the mean SSIM at each time point surveys the focality of the progression of brain states evoked by either stimuli. Comparing the SSIM between stimuli provides a metric of state colocalization. The results demonstrate that, during the first 4 to 5 seconds of the stimulus, brain states in both conditions are incoherent. After ~7 s, both conditions achieve focal brain states, with the *mentalizing* condition being much more compactly delineated. After ~12 s, both conditions assume similar brain states. The onset of the fixation block causes brain states to, again, disperse across the embedding. In the latter half of the fixation block, participants who witnessed a *mentalizing* condition tend to linger in a very tightly localized brain state. This suggests that a more characteristic variety of rumination occurs in response to the *mentalizing* stimulus. Density maps from time points having focal brain states are shown in the insets of figure 5 and close up in figure 6.

**Figure 5.**
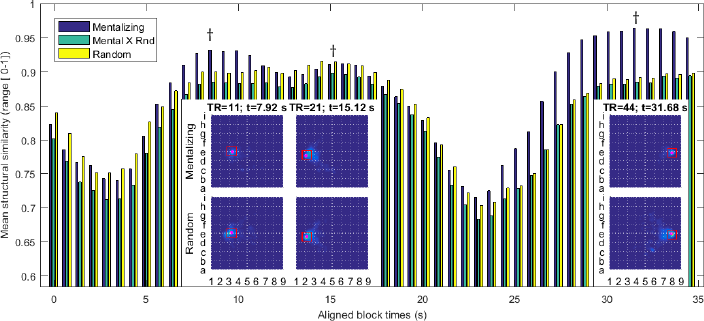
analyzes 2D Euclidean embeddings of instantaneous brain states segmented in terms of both block-design task contrasts, and by the acquisition time of temporally aligned task blocks during the SOCIAL scan. The bar chart shows the mean value of the bootstrapped structural similarity index of the within *(blue=mentalizing*, and yellow=*random*) and the between (green) block-design-contrast embeddings. For each task contrast, the sample distribution was bootstrapped from 25 realizations of 250 timepoints, randomly sampled from the group-level data set. Fewer bootstrap time points are used to accommodate the reduced sample size in each segmentation. Daggers above the bar chart point to the aligned block times whose embedding density images are displayed in the inset images. Red boxes define the boundary regions given closer scrutiny in figure 6.

**Figure 6.**
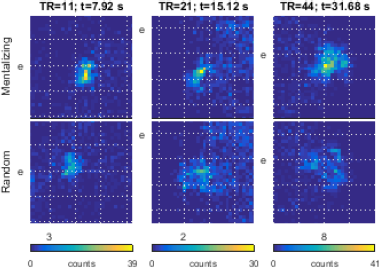
details the brain-state differences between task contrasts by displaying close-up views of the boxed areas from figure 5. Boxed areas are 1/8 the total map space, on a side. Each column is from the same map boundary region. Data are displayed as 2-dimensional histograms from a 32×32 grid inside each box. Column colorbars share the same upper limits.

To check the uniqueness of each contrast’s associated brain states, figure 6 magnifies the most densely populated regions in each contrast’s embedding at those time points presented in figure 5. After ~8 s, volunteers’ brains are observed to adopt adjacent and disjoint states. A short time later (~15 s), participants may adopt similar states albeit with the *mentalizing* stimulus inducing brains to adopt a more focal subset of the *random* stimulus’ state space. A similar observation obtains during the fixation block (~32 s) with the *mentalizing* stimuli evoking a focal subset relative to the *random* stimuli’s state space.

The extremely focused distribution of brain states during particular moments of block-design tasks motivated a closer investigation of the statistical distribution of embedded points at particular moments in time. To conduct this analysis, we generated a watershed segmentation of the embedded space after convolving all points on the embedded map with a very narrow Gaussian filter (figure 7, part A). We then labeled each embedded point in terms of the experimental condition under which the brain-state was generated, as well as in terms of the time that state was generated relative to the start of each experimental block. Next, we randomly permuted the point labels 100 times to generate a null distribution of the embedded point locations for each condition, at each time point. Finally, we calculated the z-statistic of the probability that the number of embedded points in each watershed region was greater in the real data than in the permuted data. The significance threshold was initially set to a p-value of 5%. With Bonferroni correction for multiple comparisons across ~5000 watershed regions and ~1000 individual time points, the significance threshold was set to a p-values less than 1e-8. Part B of figure 7 color-codes fine-grained watershed regions in terms of the most probable state associated with that region. For simplicity, all time points from a given condition share the same color coding. Part C of the figure addresses the inference from figures 5 and 6 that the contrasting conditions in the SOCIAL task result in highly stereotypical brain states at especially t ≅ 8s after the start of the block. As inferred from figure 6, the *social* stimulus induces highly focal brain states in the map space below and to the right of grid location e3 (green borders). While some brain states generated by the *random* stimulus overlap this region (yellow border), these brain states mostly lie above grid line e or to the left of grid line 3 (red borders).

**Figure 7.**
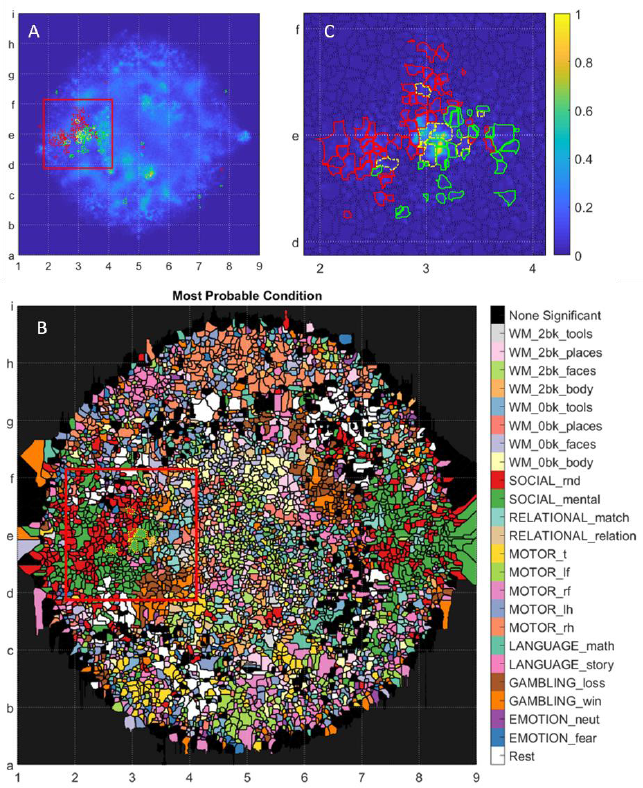
displays the statistical affinity of time-resolved and condition-dependent brain-states for a set of watershed map region. Map regions were segmented using a fine-grained density map generated from all studied time points. This granularity was motivated by the focal organization of map points in figure 6. Here, the filter width was set to 1/256^th^ the distance of the furthest map point from the map’s center. Part B displays the most probable state associated with each watershed map region. Regions where no significantly associated state was found were marked in black. Part C pursues the hypothesis that volunteers adopt different brain states around the 11^th^ image of the SOCIAL task when presented with either the *mentalizing* (green) or the *random* (red) stimulus. Map regions significantly populated in response to either stimuli, at this instant, are outlined in yellow, while regions statistically populated by only one stimulus are outlined in their respective colors. Part C’s density map is from only SOCIAL scan data. The red boxes in parts A and B outline the range of part C.

We can gain insights into how the brain responds to the contrasting stimuli by projecting a local averaging of the state space (magenta points) onto a model brain surface. Figure 8 displays the contrast between the *mentalizing* and *random* stimuli at 7.92 s (TR = 11). Whereas higher frequency (>=0.037 Hz) activations are similar, the lower frequency (<=0.019 Hz) brain states bear marked differences. At infra-slow frequencies (0.01 Hz) the *mentalizing* stimuli induces in-phase oscillations between the visual, parietal, sensorimotor, and lateral prefrontal cortices. This contrasts with brains experiencing the *random* stimulus for which the visual and left parietal cortex are out-of-phase relative to the anterior prefrontal, left orbitofrontal, and left parietal networks. Slow (0.019 Hz) activations are similar between block contrasts save for, 1) the inclusion of the medial prefrontal cortex within the positive-phase network during the *mentalizing* condition, and 2) stronger negative-phase activation in the anterior prefrontal cortex among volunteers receiving the *random* stimulus. Task-based activation studies of the same task-contrasts identified similar areas of contrasting brain activity, including the medial prefrontal cortex, lateral parietal cortices, and the visual cortex (Barch et al., 2013; Castelli et al., 2000).

**Figure 8.**
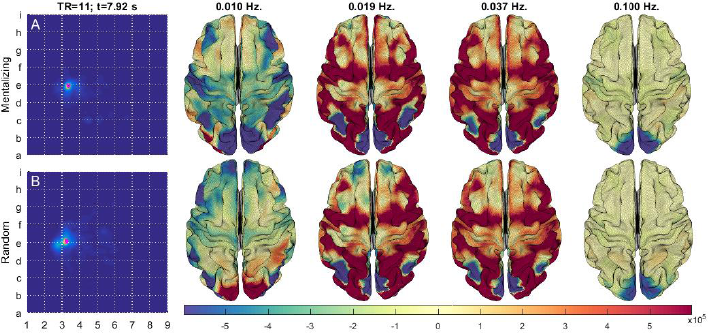
displays the difference between the brain states of participants at ~8 seconds after being given either a *mentalizing* (A) or the *random* (B) visual stimulus during the SOCIAL scan. The brain state is an average from the maximally populated embedding region from within a 32×32 grid (magenta points).

## 4. Discussion

The present analytical framework, where BOLD dynamics are interpreted as multiscalar, instantaneous events overcomes many of the challenges faced in the study of brain dynamics. Unlike methods based on sliding window correlation, it avoids the challenges involved in choosing a window length (Hindriks et al., 2016; Shakil et al., 2016). Additionally, whereas previous studies using clustering tended to delineate brain states into a fixed number, *k, of* categories (Calhoun et al., 2014), manifold embedding optimizes a low-dimensional representation of the high-dimensional data, allowing a continuous distribution. Nevertheless, it remains possible to identify discrete state categories via subsequent analyses (e.g., through a watershed transformation of the continuous state space).

The statistically significant differences between the state-space distributions of each task provides assurance that the embedded state space effectively differentiates between activation patterns related to different tasks (figures 3 and 4). On the other hand, the statistically insignificant differences between the state-space distributions of the repeated resting-state scans provide assurance that the embedded state space does not overspecify differences between brain states. The identification of fine-grained differences between the brain states of task contrasts confirms our hypothesis that common stimuli result in short distances between embedding points. Our qualitative and quantitative analysis shows that the random and mentalizing portions of the social task inhabit adjacent but disjoint map regions (figures 5 through 8).

The segmentation of the embedded space allows the identification of networks whose coactivations (or lack there-of), at particular frequencies and phases, are predominant in any given state (figure 2). Our analysis of the SOCIAL scan demonstrates this point when brain regions involved in attention (lateral prefrontal) mental representation (parietal cortex) and somatic representation (somatomotor cortex) are slowly driven, in-phase, with activations of the visual system.

One of the advantages of the 2D representation obtained from t-SNE is the ease of exploratory data analysis. From the distribution of the data, one can hypothesize about the similarities of different tasks, identify common trajectories, identify states and substates, etc. The resulting hypotheses can be addressed through further statistical analysis as demonstrated for the social task, or they might motivate more specific experiments designed to address the questions in other ways. Regardless of the following analysis, t-SNE provides a powerful tool for characterizing functional neuroimaging data.

*Insight into resting state fMRI*. Given the success of t-SNE at differentiating between tasks, the present study’s embedded state space may offer new insights into lingering questions on the character of the resting state (Lowe, 2012). The wide spatial extent, absence of low velocity regions, and overall short dwell times, converge on the finding that the resting-state is not a singular condition. As the only difference between rest and task is the absence of an explicit stimuli, the preference resting brains display for peripheral map regions marks the resting-state as mostly distinct from each of the 7 task states. One notable exception is the interior map region roughly bounded by grid vertices f5, f7, e5 and e7. Details of the region’s contribution from each task contrast (supplemental figure 5) the LANGUAGE scan’s *story* condition, the EMOTION scan’s *neutral* stimulus, WM *0 back* challenges, and the collection of time points when no stimulus information was explicitly provided (labeled rest). The region may therefore relate to times when volunteers are externally oriented albeit with low cognitive demands. Indeed, volunteer brains often populate this region during the fixation blocks of most tasks (see supplemental videos of especially the GAMBLING and RELATIONAL scans). Another notable exception is the projection of MOTOR and SOCIAL brain states to locations further out in the map’s periphery than the significant REST regions in figure 7 part B. Despite the general lack of overlap between the REST and TASK conditions, the REST condition itself exhibits several dense concentrations of time points similar to those observed during the tasks, suggesting that these network configurations constitute common brain states during the resting condition.

> Limitations. The generation of the embedding’s features comes directly from a combination of the input data and the analytical model. As always, the BOLD signal’s lack of direct sensitivity to neural activity limits our ability to infer the underlying neurophysiology from the functional imaging data. Noise in the BOLD signal (from residual physiological noise, motion, or the scanner) will affect the embedding of the data.
>
> Regarding the analysis itself, the symmetric distribution of contrasting brain-states across the map may owe itself to the use of signed wavelet coefficients. Like all spectral decompositions, the wavelet transform inherently generates phase information as complex coefficients. The present study projects complex coefficients onto the set of real numbers, thus limiting the analysis to account for two phases, separated by 180°. Other studies have found good segmentation when comparing BOLD signals across additional phases (Chang and Glover, 2010; Yaesoubi et al., 2015). The utility of incorporating phase information supports the notion that regional activations bear some degree of phase-coupling (Thompson GJ, 2014; Tort et al., 2010). Future studies may appropriate this natural feature of brain activity by characterizing the data in alternative metric spaces that better utilize complex-valued data in the high-dimensional space, and better distribute their states into a low-dimensional space.

Furthermore, it should be noted that while using as a wavelet kernel for the spectral transform is expected to result in improved time frequency localization relative to a short-time Fourier transform, it is possible to tune the kernel function to extract additional information about BOLD dynamics. In addition to requiring that the transform kernel decay smoothly to 0 after a short time, we might also require that the kernel somehow better represent the BOLD signal’s underlying impulse waveform. Such kernels may be developed by directly “lifting” the temporal shape from the data itself (Sweldens, 1998). Such a procedure would provide better localization of the BOLD signal’s energy into fewer wavelet coefficients, and thus improve the ability to differentiate between states.

*Future directions*. This study demonstrates an analytical technique to observe BOLD dynamics that performs well in segmenting contrasting activities. This approach provides a simple way to summarize patterns of brain state dynamics across various tasks as well as when participants are at rest. The ready capacity to chart brain-state dynamics against experimental stimuli raises the interest for demarcating the functional space of other varieties of conditions. Indeed, study methods are readily amenable to describing brain state dynamics of differing populations and animal models including patient populations. One potential application for the t-SNE embedding is to determine whether it can identify specific states that are present in patients but not in healthy controls. Another area of interest utilizes the preserved timing information to determine common trajectories of brain activity across states during task and rest. The t-SNE embedding facilitates exploratory analysis but can also be used to identify significant differences between tasks or populations using additional analysis. While permutation was applied for most of the statistical analyses shown in this manuscript, more sophisticated approaches should also be pursued.

The subsampling procedure reduces the computational complexity of fitting an out of sample point from O(n^2) operations to O(n). This feature enables future research to chart increasingly detailed and comprehensive maps of the brain’s dynamical state space from an ever-increasing pool of shared data. One future area of investigation should examine whether data acquired on different scanners or with different parameters (TR, for example) can be added to the existing embedding or should be handled separately.

## 5. conculsion

The BOLD signal’s multispectral components, developed naturally among the brain’s many networks, constitute a robust descriptor of instantaneous brain states. High-dimensional datasets containing point-to-point brain state dynamics are made interpretable by embedding the graph onto a 2 dimensional sheet. Our analysis of a dynamical brain-state embedding from a large population (N=446) concludes that the resting brain actively pursues a range of distinctive states from those adopted during explicit tasks. The realization of both resting and task-active states involves large-scale, and often phase-locked coordination’s among multiple brain regions at particular frequencies. The tools described in this manuscript should enable further analysis of the brain’s dynamic trajectories during both tasks and rest.

## 6. Acknowledgements

The authors gratefully acknowledge Dr. Dieter Jaeger and the Emory Center for Mind, Brain, and Culture whose *Dimensionality Reduction Workshop* inspired this work. Special thanks go to Dr. Ying Guo who suggested the permutation test for statistical significance. These efforts were directly funded by: NIH 5-R01NS078095-02, and by Professional Development Supports Funds provided by the Laney Graduate School at Emory University. Data were provided by the Human Connectome Project, WU-Minn Consortium (Principal Investigators: David Van Essen and Kamil Ugurbil; 1U54MH091657) funded by the 16 NIH Institutes and Centers that support the NIH Blueprint for Neuroscience Research; and by the McDonnell Center for Systems Neuroscience at Washington University.

## Supplemental Information

### Supplemental Methods

#### Data Acquisition

The data for this study was obtained by leveraging the library of resting-state and task fMRI images from the Human Connectome Project (HCP), a joint project between Washington University and the University of Minnesota (Van Essen et al., 2012b). These data were acquired using a customized Siemens 3T “Connectome Skyra” and the 32 channel, anterior/posterior, head receive coil. T1 weighted anatomical scans were acquired via a 3D MPRAGE sequence with *TR* = 2400 ms, *TE* = 2.14 ms, *TI* = 1000 ms, flip angle of 8°, *FOV* = 224×224 mm, and voxel size 0.7 mm isotropic. BOLD-weighted fMRI images were acquired via a gradient-echo EPI sequence with *TR = 720* ms, *TE* = 33.1 ms, flip angle of 52°, *FOV* = 208×180 mm, 72 slices, 2.0 mm isotropic voxels, and multiband factor of 8. Functional scans imaged individuals while they adopted a comprehensive battery of states. These states may be subdivided into the 9 scans named as follows: REST1, REST2, EMOTION, GAMBLING, LANGUAGE, MOTOR, RELATIONAL, SOCIAL, and WORKING MEMORY (WM). Each scan was performed twice for each volunteer, each time with an opposite phase encoding gradient (left to right, vs right to left). In total, each individual contributed 8,680 temporal and 91,282 spatial data points. REST scans spanned 4,800 time points. All data were de-identified before download.

#### Data Preprocessing

Data preprocessing include spatial artifact and distortion removal, surface generation, anatomical registration, and alignment to grayordinate space (gray-matter vertices or voxels). Subsequent use of spatial filters from a separate Independent Component Analysis (ICA) assume that data follow an isotropic noise model; thus, all voxel time series are normalized to zero mean and unit variance. To reduce the influence of edge effects during spectral filtering, contiguous, 300 image segments from a volunteer’s REST scans were placed in-between their task scans. The remaining 900 REST images capped the beginning and the end of the concatenated series with 450 time points each. The order of concatenated rest and task images were randomized across volunteers.

#### Spectral and Spatial Filtering

The BOLD signal bears a log linear relationship between power spectrum and frequency: log *S(f) = c + γlog f*; alternatively, *S(f)~1/f ^γ^*. For the average BOLD signal in brains, the power law exponent, *γ ≅ – 1*. The variable *c* is a constant. Such ’1/f-type’ systems denote that the system’s high-frequency realizations establish and maintain its low-frequency structure (Wornell, 1993). The simplest 1/f-type systems are termed,’scale-free,’ that is, one observes rescaled versions of some elementary process, a “fractal”, at all observable scales. On the other hand, complex 1/f-type systems exhibit emergent properties at multiple scales (Ciuciu et al., 2012; He, 2014; Liu et al., 2014). A theoretically optimal method for observing 1/f-type processes is to transform them using a scale-free, or multiresolution, basis set (Bullmore et al., 2004; Ciuciu et al., 2012). Coefficient variance in the scale-free domain is thus a representation of the emergence of novel signal characteristics resolved to one or more scales (Daubechies, 1992). Wavelet transforms are especially useful multiresolution transforms as their kernel functions reduce to zero over a finite time-span. By convolving a 1/f-type signal with a finite, scale-free kernel, wavelet transforms highlight the signal’s dynamical properties in both the temporal and the spectral domains.

Previous studies demonstrated that BOLD data segment into static FC subnetworks from the application of multiscale filter banks (Billings et al., 2015). In the present study, spectral filtering utilized the continuous wavelet transform with a Daubechies 4 wavelet. This continuous wavelet filterbank segmented BOLD signals into an octave of 8 frequency bands log-spaced across the decade [0.01, 0.1] Hz. This frequency range corresponds to the low-frequency fluctuation range in which BOLD fluctuations bear maximal information about neuronal activity. The mother wavelet, Daubechies 4-tap, was chosen to achieve a relatively short temporal window over each spectral band, while the number of bands is sufficient to capture the inter scale network variation observed in Billings et al. 2015.

Spatial filtering utilized a 50-component ICA decomposition. The ICA transform matric was calculated as part of the HCP beta-release of group-ICA maps (Human Connectome Project, 2014). The number of components was chosen by identifying the intersection between the eigenvalues of a volunteer’s real concatenated input data matrix, and a randomly shuffled version of that matrix, and choosing a number of components that just exceeded this point of intersect (data not shown).

#### Manifold Embedding

Each temporal sample for each volunteer’s high-dimensional state descriptor (50 spatial components by 8 spectral components) was pairwise compared using the Pearson correlation distance,

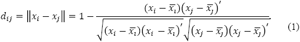

Because of the theoretically optimal whitening properties of the wavelet transform, and because we have normalized time series via z-scoring, the Pearson correlation distance highlights coordinated deviations from normative spectral intensities.

Manifold embedding was performed with the algorithm t-Distributed Stochastic Neighborhood Embedding (t-SNE) (Berman et al., 2014; van der Maaten and Hinton, 2008; van der Maaten et al., 2009). The algorithm begins by transforming high-dimensional pairwise distances into conditional probabilities, *p_j|i_*, along a Gaussian probability distribution,

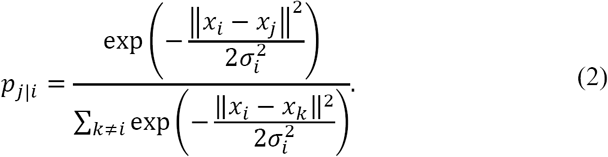

The variable *σ_i_* is equal to the variance of the high-dimensional data when multiplied by a Gaussian centered over point i. The width of each Gaussian is adjusted to cover an equivalent amount of points. Formally, the width is adjusted until the base 2 exponent of the Shannon entropy measured in the stochastic distribution around the i^th^ point achieves a fixed value termed the perplexity. For the present study, we follow the recommendation from van der Maaten & Hinton (2008) of a perplexity equal to 30. Collectively, the transformation from inter-sample distances to conditional probabilities emphasizes the natural associations of each sample to its neighbors. The authors of t-SNE also described a problem with previous implementations of SNE-based algorithms wherein moderately dissimilar samples, in the high-dimensional space, crowd together in the low-dimensional map (van der Maaten and Hinton, 2008). Therefore, t-SNE calculates the low-dimensional probabilities, *Q*, using a distribution having a much longer tail than in the high-dimensional case. A good choice to avoid this problem was found to be the Student t-distribution with one degree of freedom:

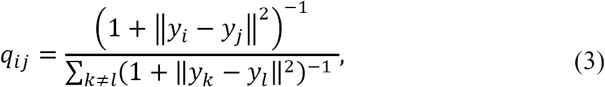

where ‖y_i_ – y_j_‖ is the Euclidean distance between samples i and j in the low-dimensional space.

A natural cost function, *C*, to calculate the fidelity of the low-dimensional representation relative to the high-dimensional data is the Kullback-Liebler (KL) divergence which is related to the cross-entropy between the two distributions. A symmetrized version of the KL divergence is used here to expedite computation time and to balance the cost of representing points that are close together in the high-dimensional space as distant points in the low-dimensional space, and vice-versa. Thus,

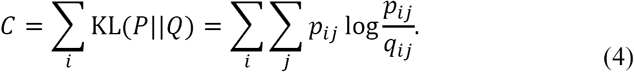

The joint probabilities in the high-dimensional space are calculated as 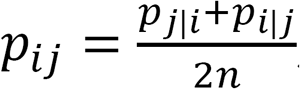, where *n* are the number of samples.

This collective description of high-dimensional and low-dimensional spaces, as well as the relationship between them, emphasizes both that similar map points are modeled by small pairwise distances and that dissimilar map points are modeled by large pairwise distances. This is the case at all but the finest scales, at which point, the numerator of equation (3) is dominated by a constant rather than by variations from the input data. The t-SNE algorithm is implemented as a gradient descent process. The form of the gradient, as well as detailed notes on methods to improve the speed of convergence may be found in van der Maaten and Hinton, 2008.

The initial construction of a t-SNE embedding is computationally expensive: O(n^2^). For a compute node having 256 GB of RAM, the maximum number of double precision data points that may be included in a single t-SNE embedding is limited to about 90,000 samples. The full complement of 4 resting state scans and 14 task scans contains 8,680 images for each of the 446 included volunteers. To overcome the computational limits of embedding larger datasets, the present study follows the recommendations from Berman et al. for training a low-dimensional embedding space from a subsampling of data points (Berman et al., 2014). Briefly, t-SNE embeddings were generated from each of 446 volunteers, individually. Next, 200 sample points were pulled from each volunteer’s embedding, at random, and in proportion to the density of points within the embedding. A group-level embedding was then trained from each volunteer’s sample of 200 time points. The best low-dimensional locations of the remaining time points vis-á-vis the trained embedding were then calculated in two steps: 1) Approximate the out-of-sample point’s low dimensional location as a weighted sum of its nearest neighbors in the full high-dimensional space. 2) Determine the local KL divergence minimum by changing only the location of the out-of-sample point. As this minimization is not convex, it is worthwhile to jitter the out-of-sample point’s initial low-dimensional location by sampling from a range of its high-dimensional neighbors. This procedure reduces the computational load to O(n). The subsampling procedure greatly increases the interpretability of the resulting map by removing the bias experienced among sequentially sampled—and hence, temporally correlated—points when they are embedded simultaneously.

#### Sub-Space Identification and Characterization

One method to summarize 2-dimensional point distributions is by convolution with a Gaussian filter. In order to account for both coarse and fine features of the embedded distribution, two filter radiuses were selected for the present study—one at 1/32 the maximum displacement from the map center and the other at 1/256. Particularly dense map regions are segmented from one another, in a data-driven fashion, by taking the watershed transform of the inverse of each density map (Meyer, 1994).

#### Velocity Field

Instantaneous velocities, were calculated by taking the difference in the embedded location of successive sample points. The group-level displacement magnitude was averaged, separately, in each of the 4 cardinal Euclidean directions, -i, +j, +i, and -j, for each point in a 32x32 grid. Results were normalized to unit magnitude.

#### Comparing Embeddings

Embeddings were segmenting against the HCP’s experimentally defined states, i.e. the resting-state and the task-based scans. To test the inference that scan-segmented maps depicted distinct brain-state distributions, we conducted an ANOVA with multiple comparisons testing using a bootstrapped sample of each experimentally defined state. Points within each bootstrap realization were chosen from segmented group-level datasets. The lower bound to the number of points in each bootstrap realization sample was chosen to ensure a full coverage of the state’s embedded range. The upper bound was chosen to ensure that few points were sampled twice in any two bootstrap realizations.

Bootstrap realizations were pairwise compared using the Structural Similarity Index (SSIM) (Zhou et al., 2004). SSIM measures the similarity between two images, *x* and *y*, as the multiplicative combination of three image quantities, the cross-luminance, I, cross-variance, *c*, and cross-structure, *s*. Thus:

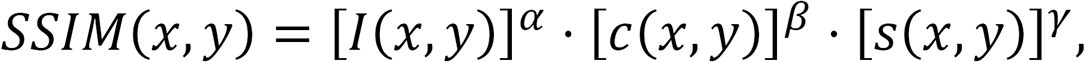

where

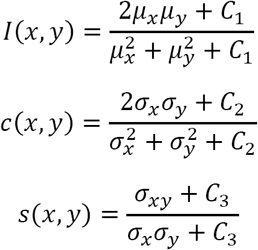

where μ_x_, μ_y_, σ_x_,σ_y_, and σ_xy_ are the local means, standard deviations, and cross-covariance for images *x, y*.The values *C_3_*, *C_3_*, and *C_3_* are small constants given by *C_1_ = (K_1_-L)^2^, C_2_ = (K_2_L*)^2^, and *C_3_* = *C_2_/2*. Here L is the dynamic range of pixel values. The variable *K_t_* ≪ 1, and the variable *K_2_* ≪ 1. The exponents over each SSIM term were set to 1 so as to weight each term equally. SSIM values range between 0, no image similarity, and 1, complete image similarity. SSIM statistical testing was conducted simultaneously for all SSIM pairs (50*50/2 independent comparisons). Maps were deemed to provide insignificant segmentation if the 95% confidence interval of the within-state SSIM fell within or below the range of any of its between-state SSIM 95% confidence intervals. The multivariate construction of the SSIM algorithm makes it a useful technique for quantifying the differences between density maps. Density maps contained equal numbers of points to ensure that the SSIM metric to remain balanced.

#### Real-Time Dynamics

Group level brain-state dynamics are characterized through map segmentation at the level of each task’s block-design contrasts. For instance, MOTION task blocks are segmented into movement of the tongue, the left hand, right hands, etc. The total set of block-design contrasts, from all individuals and each individual’s task repetitions, are aligned at time t = 0s, the start of the block (including the cue, if present). Group level density images are then calculated for each aligned image acquisition. The resting state was aligned to a single time point.

#### Permutation Testing for Labeling the Embedded map

To test the preference of labeled times and conditions for particular map regions, a null distribution was constructed by randomly permuting the labels assigned to each embedded point. Thereafter, it is possible to compare the mean number of points randomly assigned to each region, under a particular condition/time, with the actual number of points in that region, at that same condition/time. Map regions may then be labeled in terms of the preference of each region for particular condition/times.

#### Supplemental Figures

**Supplemental figure S1.**
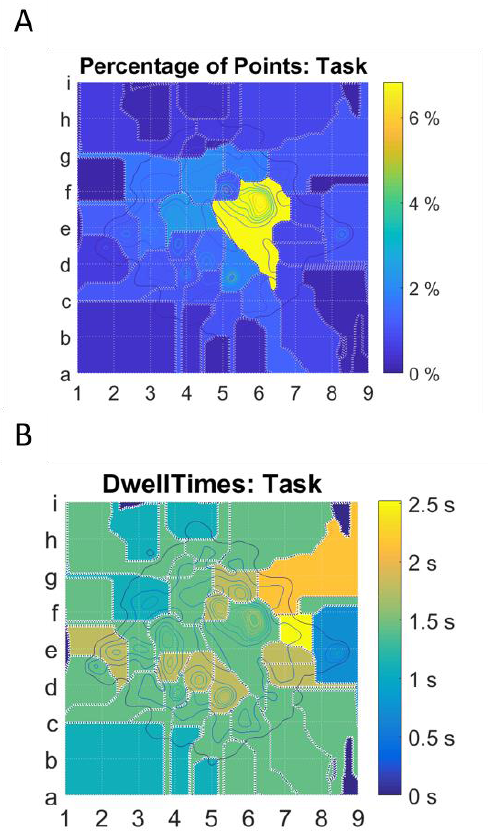
elaborates on the point distributions within watershed catchment-basins for task-active maps. Part A displays the percentage of points contained within each region. Part B displays the dwell-time for each region, reported as the mean number of temporally contiguous points contained within each region.

**Supplemental figure S2.**
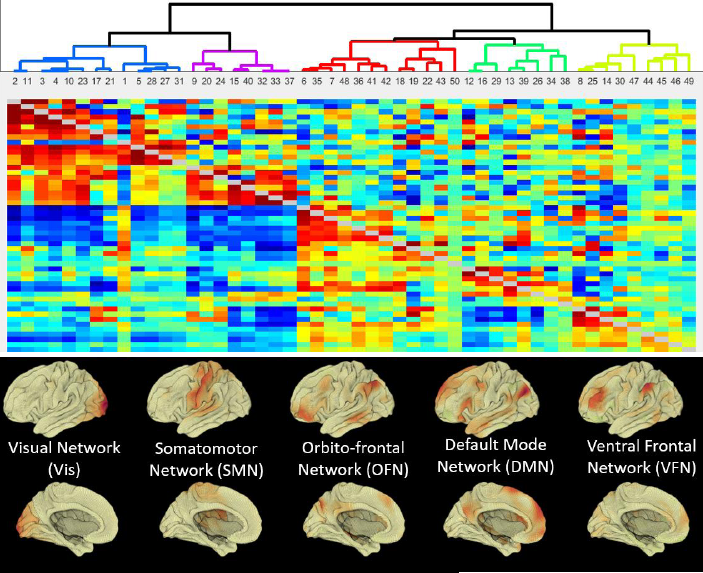
displays the relatedness between each of the 50 ICA components. Data were generated using the FSLNets toolbox provided through the HCP. The hierarchical clustering map was calculated from time-series from each ICA network back-projected for each volunteer included in the original analysis. The projection onto the brain of each of 5 ICA clusters is also shown.

**Supplemental figure S2.**
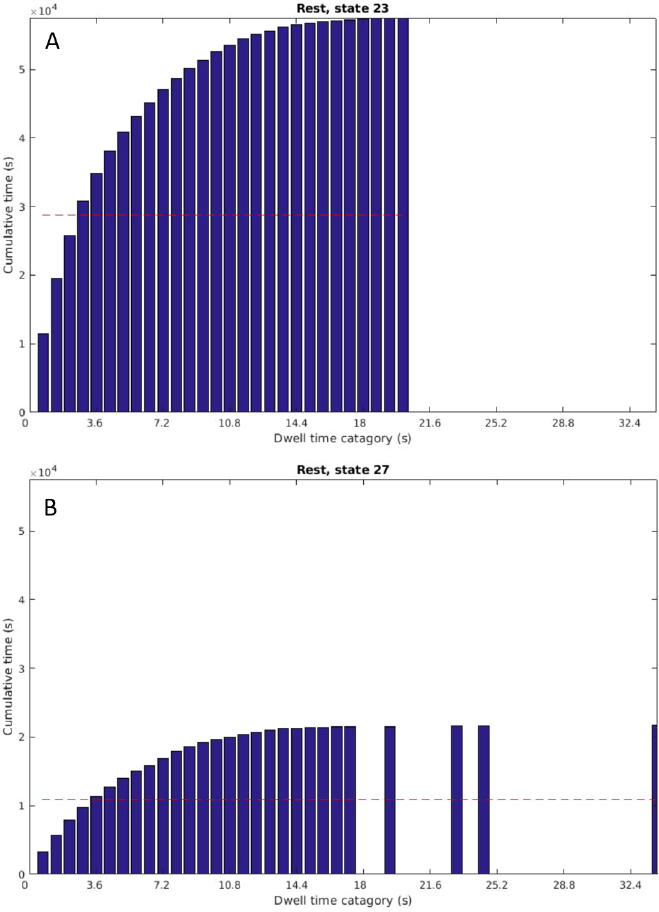
displays the dwell-time distribution for the highlighted regions in parts A and B (respectively) of figure 2.

**Supplemental figure S4.**
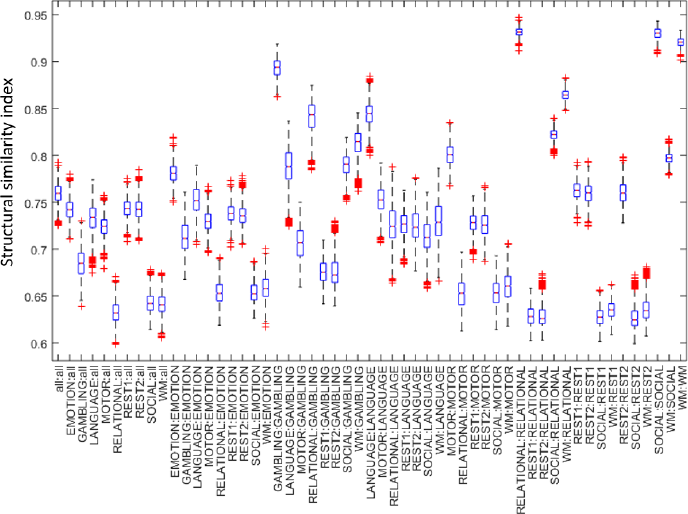
displays the descriptive statistics from bootstrap, between-scan, structural similarity index testing as a box-stem plot.

**Supplemental figure S5.**
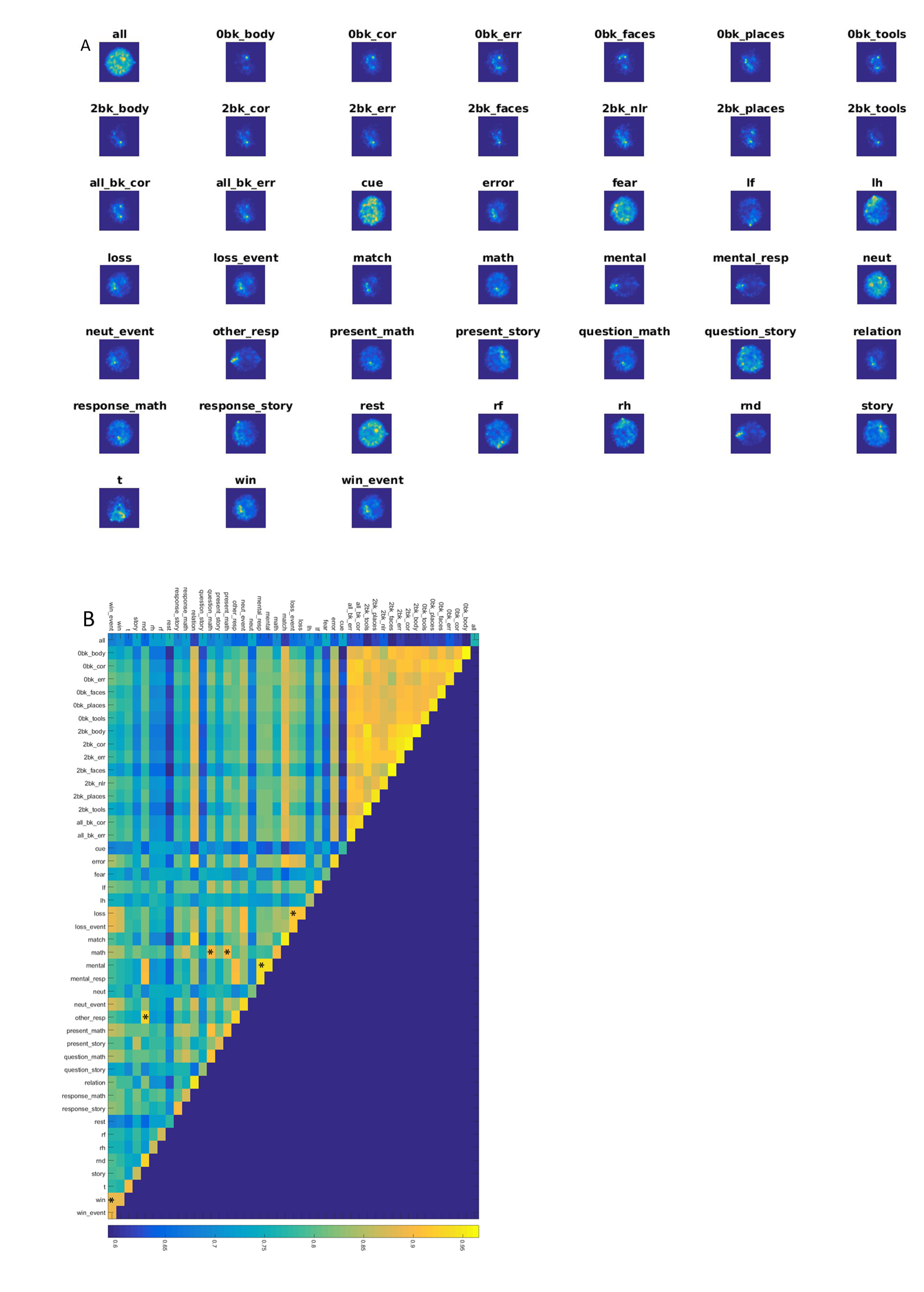
displays and compares 2D Euclidean embeddings of instantaneous brain states, segmented by within-scan events. Part A displays the density of each scan’s embedded points. Part B displays the mean structural similarity index (SSIM) from 50 bootstrap comparisons, with 2500 points per comparison. Asterisks indicate between-group comparisons whose SSIM was not significantly less than either within-group comparison.

**supplemental figure S6.**
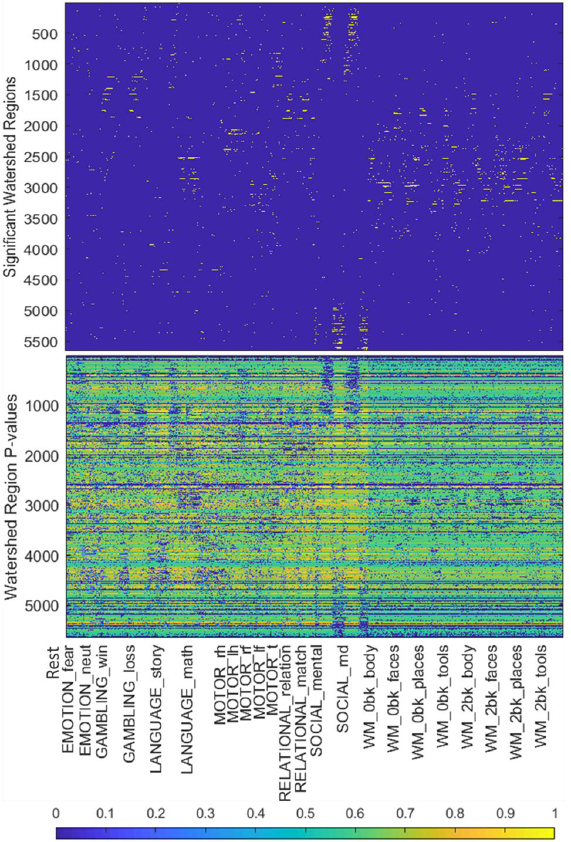
addresses the likelihood that each of the experimental condition results in any of the brain states. Conditions are aligned task blocks. The resting state is taken as a single condition. Watershed regions are from a fine-grained density map, and resulted in ~5000 regions. A z-statistic was calculated across all possible affinities. The null distribution was generated by randomly permuting the labels associated with each point 100 times. The top plot highlights statistically significant affinities (after Bonferroni correction). The bottom plot displays each comparison’s p-value.

#### Supplemental Movies

Movie 1 depicts mean brain states after segmenting the embedding using the point distribution of the resting state.

Movie 2 depicts mean brain states after segmenting the embedding using the point distribution of all tasks.

Movie 3 depicts the temporal evolution of volunteer brain states through the embedded state space during the *mental* contrast of the SOCIAL task.

Movie 4 depicts the temporal evolution of volunteer brain states through the embedded state space during the *random* contrast of the SOCIAL task.

Movie 5 depicts the temporal evolution of group-level volunteer brain states through the embedded state space during the *random* contrast of the SOCIAL task.

